# Destin2: integrative and cross-modality analysis of single-cell chromatin accessibility data

**DOI:** 10.1101/2022.11.04.515202

**Authors:** Peter Y. Guan, Jin Seok Lee, Lihao Wang, Kevin Z. Lin, Wenwen Mei, Yuchao Jiang

## Abstract

We propose Destin2, a novel statistical and computational method for cross-modality dimension reduction, clustering, and trajectory reconstruction for single-cell ATAC-seq data. The framework integrates cellular-level epigenomic profiles from peak accessibility, motif deviation score, and pseudo-gene activity and learns a shared manifold using the multimodal input, followed by clustering and/or trajectory inference. We apply Destin2 to real scATAC-seq datasets with both discretized cell types and transient cell states and carry out benchmarking studies against existing methods based on unimodal analyses. Using cell-type labels transferred with high confidence from unmatched single-cell RNA sequencing data, we adopt four performance assessment metrics and demonstrate how Destin2 corroborates and improves upon existing methods. Using single-cell RNA and ATAC multiomic data, we further exemplify how Destin’s cross-modality integrative analyses preserve true cell-cell similarities using the matched cell pairs as ground truths. Destin2 is compiled as a freely available R package available at https://github.com/yuchaojiang/Destin2.

## Introduction

Recent advances in single-cell assay of transposase-accessible chromatin followed by sequencing (scATAC-seq) technologies [1–3] offer unprecedented opportunities to characterize cellular-level chromatin accessibilities and have been successfully applied to atlas-scale datasets to yield novel insights on epigenomic heterogeneity [4, 5]. scATAC-seq data analysis presents unique methodological challenges due to its high noise, sparsity, and dimensionality [6]. Multiple statistical and computational methods have been developed and evaluated by independent benchmark studies [7].

The first set of methods call ATAC peaks or segment the genome into bins and take the cell by peak (or cell by bin) matrix as input. Among these methods, Signac [8], scOpen [9], and RA3 [10] perform TF-IDF normalization followed by different dimension reduction techniques. SnapATAC [11] computes a Jaccard similarity matrix, while cisTopic [12] performs topic modeling. Moving beyond the peak matrix, Cicero [13] and MAESTRO [14] make gene expression predictions from unweighted and weighted sum of the ATAC reads in gene bodies and promoter regions, respectively; the predicted gene activities have been shown to in the ballpark recapitulate the transcriptomic profiles and discern cell populations [15]. For TF-binding motifs, chromVAR [16] computes a motif deviation score by estimating the gain or loss of accessibility within peaks sharing the same motif relative to the average cell profile; these deviation scores have also been shown to enable accurate clustering of scATAC-seq data.

Notably, most, if not all, of the aforementioned methods carry out “unimodal” analysis with a single type of feature input (i.e., peaks, genes, or motifs). One of the earliest methods, SCRAT [17], proposes to use empirical and prior knowledge to aggregate the peaks into genes, motifs, and gene sets, while neglecting the peak-level information due to high computational burden. EpiScanpy [18], ArchR [19], and Signac [8] all generate multimodal feature inputs. However, dimension reduction and clustering are still focused on the peak accessibilities – the gene activities are generally integrated with singlecell RNA sequencing (scRNA-seq) data for alignment, and the motif deviation scores are used to identify enriched and/or differentially accessible motifs.

To our best knowledge, no integrative methods are available for a cross-modality analysis of scATAC- seq data, yet it has been shown that the peaks, genes, and motifs all contain signals to separate the different cell types/states. Here, we propose Destin2, a successor to our previous unimodal method Destin [6], for cross-modality dimension reduction, clustering, and trajectory reconstruction for scATAC-seq data. The framework integrates cellular-level epigenomic profiles from peak accessibility, motif deviation score, and pseudo-gene activity and learns a shared manifold using the multimodal input. We apply the method to real datasets with both discretized cell types and transient cell states and carry out benchmarking studies to demonstrate how Destin’s cross-modality integration corroborates and improves upon existing methods based on unimodal analyses.

## Materials and Methods

Figure 1 outlines Destin2’s analytical framework. For unimodal data input, Destin2 utilizes Signac [8], MAESTRO [14], and chromVAR [16] for pre-processing and generating the matrices of peak accessibility, gene activity, and motif deviation, where the cell dimensions are matched, and yet the feature dimensions differ. The peak matrix can be directly loaded from the output of cellranger-atac or called/refined by MACS2 [20]. Pseudo-gene activities can be derived from either taking the sum of ATAC reads in gene bodies and promoter regions by Signac [8] or using a regulatory potential model that sums ATAC reads weighted based on existing gene annotations by MAESTRO [14]. Motif deviation scores are computed using chromVAR [16] and measure the deviation in chromatin accessibility across the set of peaks containing the TF-binding motifs, compared to a set of background peaks. Destin2, by its default, uses the JASPAR database for pairs of TF and motif annotation in vertebrates [21].

**Figure 1.**
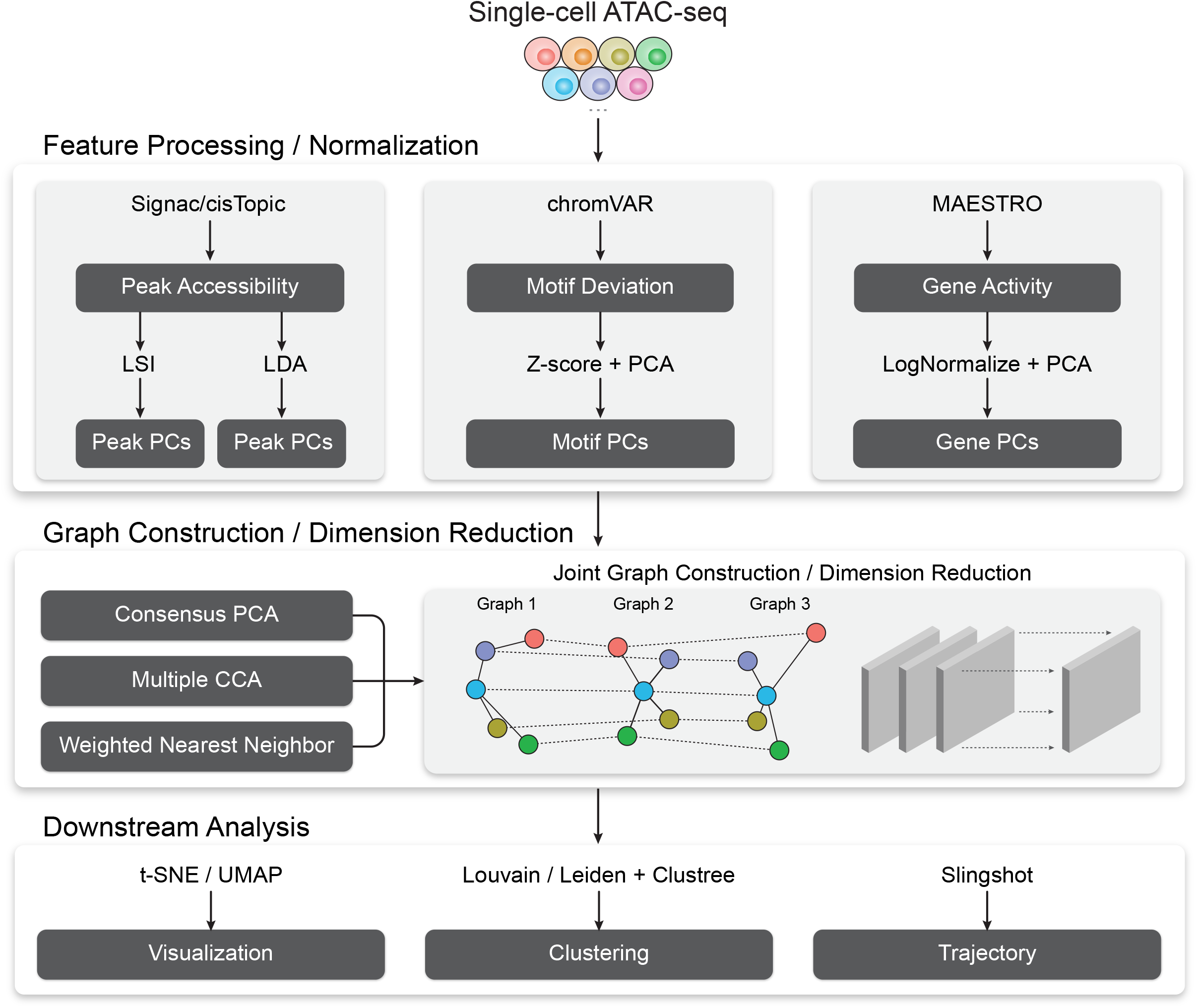
A flowchart outlining the procedures for cross-modality scATAC-seq analysis by Destin2.

For data normalization and dimension reduction, we adopt two parallel and state-of-the-art approaches, latent semantic indexing (LSI) and latent Dirichlet allocation (LDA), for the peak matrix. LSI normalizes reads within peaks using the term frequency-inverse document frequency transformation (TF-IDF), followed by a PCA-based dimension reduction [8]. LDA is a topic modeling approach commonly used in natural language processing and has been successfully applied to scATAC-seq data to identify cell states from topic-cell distribution and explore *cis*-regulatory regions from region-topic distribution by cisTopic [12]. For the motif and gene matrix, we use z-score transformation and the LogNormalize function by Seurat [22], followed by principal component analysis (PCA), respectively. These within-modality normalization and dimension reduction, which return peak principal components (PCs), motif PCs, and gene PCs, are necessary. They effectively reduce signal-to-noise ratios, and more importantly, it has been shown that PCA, followed by canonical correlation analysis (CCA), offers a powerful approach to uncover latent structure shared across modalities through an integrative analysis [23]. The number of PCs can be chosen by inspecting the variance reduction (i.e., elbow) plot or using the JackStraw method [24], which randomly permutes a subset of the data and compares the PCs for the permuted data with the observed PCs to determine statistical significance.

With the pre-processed and normalized unimodal data input, Destin2 offers three options for cross-modality integration: consensus PCA (CPCA), generalized/multiple CCA (MultiCCA), and weighted nearest neighbor (WNN). Denote the feature input across *K* modalities as *X*^(1)^ ∈ ℝ^*n*×*P*_1_^, …,*X*∈ ℝ^*n*×*p_K_*^, where the *n* cells are matched. (I) CPCA [25], algebraically equivalent to applying a second-step PCA to the concatenated peak PCs, motif PCs, and gene PCs, returns consensus PCs as joint dimension reductions, which reveal the union of the latent structure across multiple modalities. To identify the first-rank consensus PC is analogous to solve:

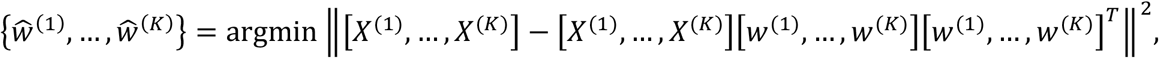

such that *w* ∈ ℝ^*p*_*k*_^ for 1 ≤ *k* ≤ *K* and 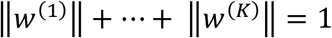. (II) MultiCCA [26], on the other hand, finds maximally correlated linear combinations of the features between each pair of modalities by solving:

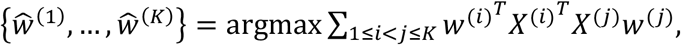

such that *w*^(*k*)^ ∈ℝ^*p*_*k*_^ and 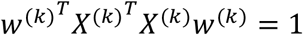 for 1 ≤ *k* ≤ *K*; we utilize the implementation from the mogsa package [27] in R. (III) Instead of optimizing for the modality- and feature-specific loading vector for projections of the three modalities, the recently developed WNN method [28] learns cell- and modality-specific weights, which reflect the information content for each modality and are used to calculate a weighted cell-cell similarity measure and construct a WNN graph. We will not go into the algorithmic details of the WNN method – readers can refer to the Seurat V4 publication [28], where the WNN framework is extended to more than two modalities with matched cells.

Followed by joint dimension reduction and graph construction, tSNE/UMAP can be used for visualization. Destin2 adopts Louvain/Leiden clustering [29] for community detection and identification of discrete cell clusters. The number of cell clusters (i.e., the resolution parameter) can be optimized using the clustree method [30], which builds a tree to visualize and examine how clusters are related to each other at varying resolutions, allowing researchers to assess which clusters are distinct and which are unstable with the use of additional metrics such as the SC3 stability index [31]. For cell population exhibiting continuous and connected cell states, Destin2 resorts to a flexible and modularized approach, Slingshot [32], for trajectory reconstruction; smooth representation of the lineages and pseudotime values are inferred using the joint dimension reduction and visualized on the UMAP space.

## Results

### Destin2 improves clustering accuracy compared to unimodal analysis methods

We apply Destin2 to four scATAC-seq datasets of human peripheral blood mononuclear cells (PBMCs) from 10x Genomics, adult mouse cortex cells from 10x Genomics, human bone marrow mononuclear cells (BMMCs) [33], and human fetal organs [5]. See Supplementary Table 1 for summary and details of the data. For the PBMC and adult mouse cortex datasets, we annotate cell types using scRNA-seq experiments from the same biological systems (PBMC from 10x Genomics and mouse brain from the Allen Brain Institute), utilizing the CCA-based method for cross-modality integration and label transfer [34] and only keeping cells that can be uniquely and confidently assigned to one cell type. For the BMMC dataset, we use the curated cell type labels from the original publication [33]. For the human fetal dataset, we resort to the tissues of origin from the experimental design/sample collection. These cell types/tissues are used as ground truths for performance assessment.

We apply unimodal analysis methods (i.e., peak analysis by Signac and cisTopic, motif analysis by chromVAR, and gene activity analysis by Signac/MAESTRO) and Destin2 to these datasets, with UMAP visualizations shown in Supplementary Figure 1. For benchmarking, we adopt four metrics for performance assessment. (I) Adjusted rand index (ARI) is used to compare the identified cell clusters against the annotated cell types, with 1 indicating that the two are exactly the same. (II) Adjusted mutual information (AMI) is similar to ARI but is more suited when there exist small and unbalanced clusters [35]. (III) Homogeneity score (H-score) is an entropy-based measure of the similarity between two clusterings and ranges between 0 and 1, where 1 indicates perfect homogeneity. (IV) Cell-type local inverse Simpson’s index (cLISI) [36] is used to assess the degree of mixing/separation of annotated cell types, with 1 indicating that the different cell types group separately and 2 indicating that the different cell types erroneously group together.

Across the four scATAC-seq datasets, our results suggest that the multimodal analysis methods proposed by Destin2 improve clustering accuracy compared to conventional unimodal analysis methods using ARI and AMI as assessment metrics (Figure 2, Supplementary Table 2). For cLISI and H-score, since the gold-standard cell-type labels are transferred using the LSI-based dimension reduction as weights, it is not surprising that the LSI method achieves the top performance; nonetheless, the difference between LSI-based methods and Destin2’s cross-modality integration results are negligible (Figure 2, Supplementary Table 2). Note that while the motif analysis returns the seemingly worst result, whether the motif modality is included in the integrative analysis does not significantly alter the output (Supplementary Figure 2), demonstrating Destin2’s robustness to the differential information content across modalities. Additionally, and more importantly, careful inspection of the confusion matrix (shown as a heatmap in Supplementary Figure 3) suggests that Destin2 is able to identify cell types/states that are otherwise indistinguishable and/or wrongly classified from a unimodal analysis – e.g., Lamp5 v.s. Vip in Supplementary Figure 3B, as well as GMP v.s. CD14 monocytes and CLP v.s. pre-B cells in Supplementary Figure 3C.

**Figure 2.**
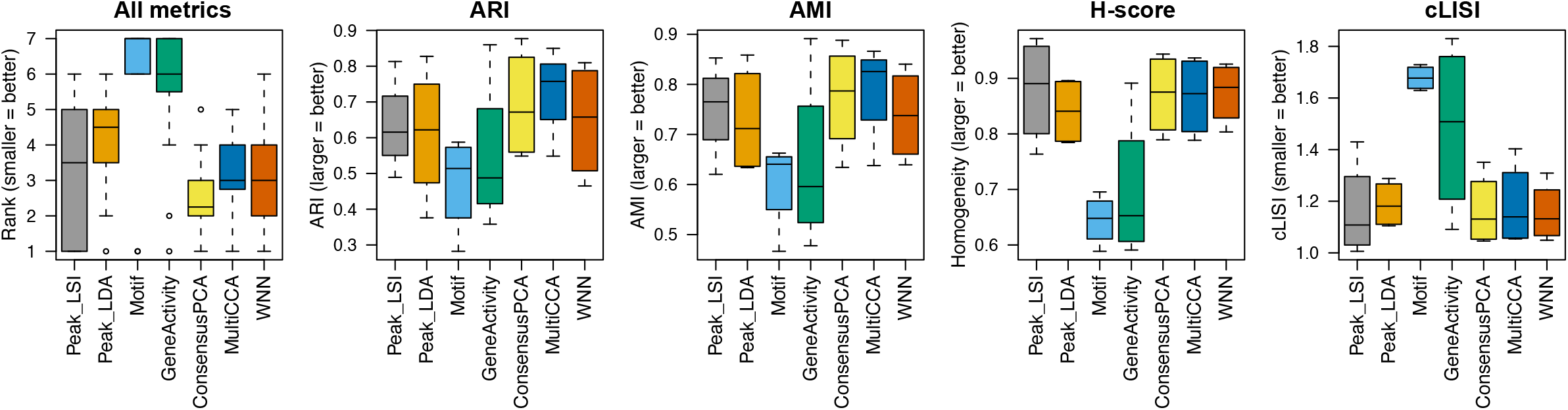
Benchmarking clustering accuracy. Four different metrics – ARI, AMI, H-score, and cLISI – were used for performance assessment. For each metric, results from the four scATAC-seq datasets are aggregated; across all metrics, the ranks of the methods are calculated. Destin2’s multimodal analysis framework achieves the highest rank and improves clustering accuracy compared to conventional unimodal analysis methods.

For downstream analysis, we first demonstrate how to determine the clustering resolution using the clustree method (Supplementary Figure 4). Specifically, clustering results with varying clustering resolutions (and thus varying SC3 stability measures [31]) are visualized as a tree: new clusters form from existing clusters, and the overlap in cells between clusters at adjacent resolutions is computed and used to calculate the in-proportion for each edge. Unstable clusters result in cells switching between branches of the trees, with low in-proportion edges; one can thus infer which areas of the tree are likely to be the result of true clusters and which are caused by over-clustering [30]. For cell populations with continuous cell states, we further demonstrate how to reconstruct the development/differentiation trajectory using Destin2’s joint dimension reduction paired with the Slingshot method. As an example, we show the reconstruction of the true branching lineages during human hematopoietic differentiation using the BMMC data (Supplementary Figure 5).

### Destin2 better preserves cell-cell similarities using single-cell RNA and ATAC multiomic data

We further applied Destin2 to three single-cell RNA and ATAC multiomic datasets of human PBMCs from 10x Genomics, adult mouse cortex cells from 10x Genomics, and mouse skin data from SHARE-seq [37]. See Supplementary Table 1 for a data summary. In using these multiomic datasets, we demonstrate how Destin2’s cross-modality analyses preserve true cell-cell similarities by using the matched cell information as ground truth without relying on methods for cross-modality alignment. Importantly, this also does not need the RNA-ATAC alignment or the transfer of discretized cell-type labels, which often fails for cell populations consisting of transient states.

We apply the same unimodal analysis methods and Destin2 to these datasets. For benchmarking, we adopt two additional metrics designed specifically for the single-cell multiomic data – fraction of samples closer than the nearest neighbor (FOSCTTNN) and agreement. Both metrics measure the preservation of a cell’s nearest neighbors between the RNA and ATAC domains and do not rely on annotated cell types or identified clusters. FOSCTTNN is adapted from the “fraction of samples closer than the true match” metric [38]: for each cell, we first identify its nearest neighbor (i.e., closest cell) in the RNA domain as ground truth, and then, in the ATAC domain, we calculate the fraction of cells that are closer than its true nearest neighbor. For agreement [39], we identify each cell’s *k* nearest neighbors in the RNA and ATAC domains, respectively, and then calculate the fraction of overlap. Nearest neighbors are identified using Euclidean distance of the cells’ reduced dimension from each modality and method. The two cell-specific metrics can be further summarized using median and Gini mean difference (GMD) across cells.

For the three multiomic datasets, our results suggest that the multimodal analysis methods offered by Destin2 exhibit top or near-top performance. For FOSCTTNN, Destin2’s cross-modality integration results are either top-performing or negligibly different from the top performer (Table 1); for agreement across different *k* (number of nearest neighbors), the WNN method achieves the top performance (Table 2). Interestingly and importantly, neither LSI nor LDA is indefinitely preferred from this benchmark analysis – e.g., LDA outperforms LSI using the FOSCTTNN metric in the PBMC data (Table 1A), while LSI improves upon LDA using the agreement metric by a large margin in the mouse brain data (Table 2B). In real data analysis, where there is no ground truth to guide method selection, Destin2 integrates and corroborates information from both methods and demonstrates its robustness.

**Table 1.** FOSCTTNN metrics on single-cell RNA and ATAC multiomic datasets. Destin2’s multimodal analyses achieve top or near-top performance. FOSCTTNN is bound between 0 and 1, with 0 being the best performance. Neither LSI nor LDA is indefinitely preferred from this benchmark analysis; Destin2 integrates and corroborates information across methods and modalities.

| Method | q1 | median | GMD | mean | sd | q3 |
| --- | --- | --- | --- | --- | --- | --- |
| Peak_LSI | 0.015 | 0.045 | 0.107 | 0.089 | 0.126 | 0.116 |
| Peak_LDA | 0.013 | 0.039 | 0.081 | 0.069 | 0.095 | 0.090 |
| Motif | 0.048 | 0.131 | 0.168 | 0.174 | 0.158 | 0.261 |
| GeneActivity | 0.017 | 0.054 | 0.124 | 0.104 | 0.133 | 0.137 |
| ConsensusPCA | 0.014 | 0.045 | 0.098 | 0.084 | 0.107 | 0.118 |
| MultiCCA | 0.014 | 0.044 | 0.104 | 0.086 | 0.121 | 0.112 |
| WNN | 0.013 | 0.042 | 0.094 | 0.080 | 0.102 | 0.110 |

(B) Mouse Brain 10x Genomics FOSCTTNN
| Method | q1 | median | GMD | mean | sd | q3 |
| --- | --- | --- | --- | --- | --- | --- |
| Peak_LSI | 0.004 | 0.014 | 0.085 | 0.056 | 0.130 | 0.043 |
| Peak_LDA | 0.006 | 0.021 | 0.077 | 0.056 | 0.110 | 0.056 |
| Motif | 0.010 | 0.033 | 0.105 | 0.079 | 0.133 | 0.085 |
| GeneActivity | 0.028 | 0.109 | 0.308 | 0.257 | 0.294 | 0.487 |
| ConsensusPCA | 0.005 | 0.017 | 0.088 | 0.059 | 0.129 | 0.049 |
| MultiCCA | 0.005 | 0.017 | 0.083 | 0.057 | 0.122 | 0.050 |
| WNN | 0.005 | 0.018 | 0.091 | 0.062 | 0.130 | 0.055 |

(C) Mouse Skin SHARE-seq FOSCTTNN
| Method | q1 | median | GMD | mean | sd | q3 |
| --- | --- | --- | --- | --- | --- | --- |
| Peak_LSI | 0.011 | 0.034 | 0.165 | 0.117 | 0.196 | 0.110 |
| Peak_LDA | 0.011 | 0.036 | 0.143 | 0.104 | 0.174 | 0.101 |
| Motif | 0.024 | 0.076 | 0.185 | 0.154 | 0.194 | 0.200 |
| GeneActivity | 0.062 | 0.208 | 0.332 | 0.323 | 0.301 | 0.551 |
| ConsensusPCA | 0.011 | 0.034 | 0.147 | 0.105 | 0.180 | 0.097 |
| MultiCCA | 0.011 | 0.034 | 0.139 | 0.100 | 0.176 | 0.092 |
| WNN | 0.010 | 0.031 | 0.134 | 0.096 | 0.170 | 0.091 |

**Table 2.** Agreement metrics on single-cell RNA and ATAC multiomic datasets. Different numbers of nearest numbers (*k*) were selected. Destin2’s multimodal analyses, especially the WNN method, achieve top or near-top performance. Agreement is bound between 0 and 1, with 1 being the best performance. Neither LSI nor LDA is indefinitely preferred from this benchmark analysis; Destin2 integrates and corroborates information across methods and modalities.

(A) PBMC 10x Genomics Agreement
| k | Peak_LSI | Peak_LDA | Motif | GeneActivity | ConsensusPCA | MultiCCA | WNN |
| --- | --- | --- | --- | --- | --- | --- | --- |
| 50 | 0.091 | 0.102 | 0.032 | 0.085 | 0.092 | 0.091 | 0.107 |
| 100 | 0.163 | 0.175 | 0.061 | 0.150 | 0.163 | 0.163 | 0.183 |
| 150 | 0.220 | 0.232 | 0.086 | 0.200 | 0.219 | 0.221 | 0.241 |
| 200 | 0.269 | 0.282 | 0.110 | 0.244 | 0.268 | 0.270 | 0.290 |

(B) Mouse Brain 10x Genomics Agreement
| k | Peak_LSI | Peak_LDA | Motif | GeneActivity | ConsensusPCA | MultiCCA | WNN |
| --- | --- | --- | --- | --- | --- | --- | --- |
| 50 | 0.350 | 0.288 | 0.222 | 0.119 | 0.324 | 0.318 | 0.314 |
| 100 | 0.514 | 0.439 | 0.358 | 0.196 | 0.481 | 0.478 | 0.468 |
| 150 | 0.613 | 0.549 | 0.455 | 0.255 | 0.584 | 0.583 | 0.569 |
| 200 | 0.681 | 0.629 | 0.532 | 0.306 | 0.659 | 0.658 | 0.643 |

| k | Peak_LSI | Peak_LDA | Motif | GeneActivity | ConsensusPCA | MultiCCA | WNN |
| --- | --- | --- | --- | --- | --- | --- | --- |
| 50 | 0.046 | 0.048 | 0.024 | 0.010 | 0.046 | 0.046 | 0.050 |
| 100 | 0.088 | 0.088 | 0.047 | 0.020 | 0.087 | 0.085 | 0.094 |
| 150 | 0.124 | 0.122 | 0.068 | 0.030 | 0.123 | 0.122 | 0.131 |
| 200 | 0.157 | 0.154 | 0.087 | 0.039 | 0.156 | 0.155 | 0.166 |

## Discussion

We propose Destin2 to integrate multimodal peak accessibility, motif deviation, and pseudo-gene activity measures derived from scATAC-seq data. For peak accessibility, Destin2 integrates two most popular techniques – LSI and LDA – for data pre-processing and within-modality dimension reduction. While Destin2 is not restricted to only taking peak accessibilities as input, our framework offers a strategy to ensemble results from the various peak-modeling methods [7], so long as method-specific dimension reductions are provided. For motif deviation, Destin2, by its default, resorts to chromVAR [16]. However, chromVAR can be computationally intensive and infeasible to handle atlas-scale scATAC-seq data – additional efforts are needed for higher scalability. While alternative methods are currently being developed, a viable shortcut solution is to process the entire data in mini batches (i.e., random subsamples of the cells), thus not requiring all the data to be loaded into memory at one time. Such a strategy has been successfully applied to scRNA-seq data with millions of cells [40]. For pseudo-gene activity, Destin2 aggregates ATAC reads over gene bodies and promoter regions using Signac [8] or MAESTRO [14], yet this largely neglects peaks and fragments from intergenic and noncoding regions. Additional annotations, such as the enhancers [41], super-enhancers [42], A/B compartments [43], and chromatin loops [44], can be easily incorporated into Destin2’s framework as additional modalities to be integrated.

For cross-modality joint modeling, Destin2 utilizes three statistically rigorous and computationally efficient methods – CPCA, MultiCCA, and WNN – and shows its outperformance and robustness. Additional methods that fall in the realm of multiomic integration (e.g., JIVE [45], MOFA [46], etc.) have not been thoroughly explored. For CCA, its variants and extensions, such as sparse CCA [47] and decomposition-based CCA [48], can potentially further boost performance. Overall, we believe that the framework by Destin2 introduces the concept of multiomic integration to scATAC-seq data; through the various benchmarking studies, we exemplify its utility and benefit and illustrate how it can better facilitate downstream analyses.

## Author Contributions

Y.J. initiated and envisioned the study. P.Y.G., J.S.L., L.W., and Y.J. developed and implemented the algorithm. All authors carried out data analyses. P.Y.G. compiled the R package. P.Y.G. and Y.J. wrote the manuscript, which was edited and approved by all authors.

## References

1. Buenrostro JD, Wu B, Litzenburger UM, Ruff D, Gonzales ML, Snyder MP, Chang HY, Greenleaf WJ: Single-cell chromatin accessibility reveals principles of regulatory variation. Nature 2015, 523:486–490.

2. Cusanovich DA, Daza R, Adey A, Pliner HA, Christiansen L, Gunderson KL, Steemers FJ, Trapnell C, Shendure J: Multiplex single cell profiling of chromatin accessibility by combinatorial cellular indexing. Science 2015, 348:910–914.

3. Satpathy AT, Granja JM, Yost KE, Qi Y, Meschi F, McDermott GP, Olsen BN, Mumbach MR, Pierce SE, Corces MR, et al: Massively parallel single-cell chromatin landscapes of human immune cell development and intratumoral T cell exhaustion. Nat Biotechnol 2019, 37:925–936.

4. Cusanovich DA, Hill AJ, Aghamirzaie D, Daza RM, Pliner HA, Berletch JB, Filippova GN, Huang X, Christiansen L, DeWitt WS, et al: A Single-Cell Atlas of In Vivo Mammalian Chromatin Accessibility. Cell 2018, 174:1309–1324 e1318.

5. Domcke S, Hill AJ, Daza RM, Cao J, O’Day DR, Pliner HA, Aldinger KA, Pokholok D, Zhang F, Milbank JH, et al: A human cell atlas of fetal chromatin accessibility. Science 2020, 370.

6. Urrutia E, Chen L, Zhou H, Jiang Y: Destin: toolkit for single-cell analysis of chromatin accessibility. Bioinformatics 2019, 35:3818–3820.

7. Chen H, Lareau C, Andreani T, Vinyard ME, Garcia SP, Clement K, Andrade-Navarro MA, Buenrostro JD, Pinello L: Assessment of computational methods for the analysis of single-cell ATAC-seq data. Genome Biol 2019, 20:241.

8. Stuart T, Srivastava A, Madad S, Lareau CA, Satija R: Single-cell chromatin state analysis with Signac. Nat Methods 2021, 18:1333–1341.

9. Li Z, Kuppe C, Ziegler S, Cheng M, Kabgani N, Menzel S, Zenke M, Kramann R, Costa IG: Chromatin-accessibility estimation from single-cell ATAC-seq data with scOpen. Nat Commun 2021, 12:6386.

10. Chen S, Yan G, Zhang W, Li J, Jiang R, Lin Z: RA3 is a reference-guided approach for epigenetic characterization of single cells. Nat Commun 2021, 12:2177.

11. Fang R, Preissl S, Li Y, Hou X, Lucero J, Wang X, Motamedi A, Shiau AK, Zhou X, Xie F, et al: Comprehensive analysis of single cell ATAC-seq data with SnapATAC. Nat Commun 2021, 12:1337.

12. Bravo Gonzalez-Blas C, Minnoye L, Papasokrati D, Aibar S, Hulselmans G, Christiaens V, Davie K, Wouters J, Aerts S: cisTopic: cis-regulatory topic modeling on single-cell ATAC-seq data. Nat Methods 2019, 16:397–400.

13. Pliner HA, Packer JS, McFaline-Figueroa JL, Cusanovich DA, Daza RM, Aghamirzaie D, Srivatsan S, Qiu X, Jackson D, Minkina A, et al: Cicero Predicts cis-Regulatory DNA Interactions from Single-Cell Chromatin Accessibility Data. Mol Cell 2018, 71:858–871 e858.

14. Wang C, Sun D, Huang X, Wan C, Li Z, Han Y, Qin Q, Fan J, Qiu X, Xie Y, et al: Integrative analyses of single-cell transcriptome and regulome using MAESTRO. Genome Biol 2020, 21:198.

15. Jiang Y, Harigaya Y, Zhang Z, Zhang H, Zang C, Zhang NR: Nonparametric single-cell multiomic characterization of trio relationships between transcription factors, target genes, and cis-regulatory regions. Cell Syst 2022, 13:737–751 e734.

16. Schep AN, Wu B, Buenrostro JD, Greenleaf WJ: chromVAR: inferring transcription-factor-associated accessibility from single-cell epigenomic data. Nat Methods 2017, 14:975–978.

17. Ji Z, Zhou W, Ji H: Single-cell regulome data analysis by SCRAT. Bioinformatics 2017, 33:2930–2932.

18. Danese A, Richter ML, Chaichoompu K, Fischer DS, Theis FJ, Colome-Tatche M: EpiScanpy: integrated single-cell epigenomic analysis. Nat Commun 2021, 12:5228.

19. Granja JM, Corces MR, Pierce SE, Bagdatli ST, Choudhry H, Chang HY, Greenleaf WJ: ArchR is a scalable software package for integrative single-cell chromatin accessibility analysis. Nat Genet 2021, 53:403–411.

20. Zhang Y, Liu T, Meyer CA, Eeckhoute J, Johnson DS, Bernstein BE, Nusbaum C, Myers RM, Brown M, Li W, Liu XS: Model-based analysis of ChIP-Seq (MACS). Genome Biol 2008, 9:R137.

21. Fornes O, Castro-Mondragon JA, Khan A, van der Lee R, Zhang X, Richmond PA, Modi BP, Correard S, Gheorghe M, Baranasic D, et al: JASPAR 2020: update of the open-access database of transcription factor binding profiles. Nucleic Acids Res 2020, 48:D87–D92.

22. Butler A, Hoffman P, Smibert P, Papalexi E, Satija R: Integrating single-cell transcriptomic data across different conditions, technologies, and species. Nat Biotechnol 2018, 36:411–420.

23. Brown BC, Bray NL, Pachter L: Expression reflects population structure. PLoS Genet 2018, 14:e1007841.

24. Satija R, Farrell JA, Gennert D, Schier AF, Regev A: Spatial reconstruction of single-cell gene expression data. Nat Biotechnol 2015, 33:495–502.

25. Westerhuis JA, Kourti T, MacGregor JF: Analysis of multiblock and hierarchical PCA and PLS models. Journal of Chemometrics: A Journal of the Chemometrics Society 1998, 12:301–321.

26. Kettenring JR: Canonical analysis of several sets of variables. Biometrika 1971, 58:433–451.

27. Meng C, Basunia A, Peters B, Gholami AM, Kuster B, Culhane AC: MOGSA: Integrative Single Sample Gene-set Analysis of Multiple Omics Data. Mol Cell Proteomics 2019, 18:S153–S168.

28. Hao Y, Hao S, Andersen-Nissen E, Mauck WM, 3rd, Zheng S, Butler A, Lee MJ, Wilk AJ, Darby C, Zager M, et al: Integrated analysis of multimodal single-cell data. Cell 2021, 184:3573–3587 e3529.

29. Traag VA, Waltman L, van Eck NJ: From Louvain to Leiden: guaranteeing well-connected communities. Sci Rep 2019, 9:5233.

30. Zappia L, Oshlack A: Clustering trees: a visualization for evaluating clusterings at multiple resolutions. Gigascience 2018, 7.

31. Kiselev VY, Kirschner K, Schaub MT, Andrews T, Yiu A, Chandra T, Natarajan KN, Reik W, Barahona M, Green AR, Hemberg M: SC3: consensus clustering of single-cell RNA-seq data. Nat Methods 2017, 14:483–486.

32. Street K, Risso D, Fletcher RB, Das D, Ngai J, Yosef N, Purdom E, Dudoit S: Slingshot: cell lineage and pseudotime inference for single-cell transcriptomics. BMC Genomics 2018, 19:477.

33. Granja JM, Klemm S, McGinnis LM, Kathiria AS, Mezger A, Corces MR, Parks B, Gars E, Liedtke M, Zheng GXY, et al: Single-cell multiomic analysis identifies regulatory programs in mixed-phenotype acute leukemia. Nat Biotechnol 2019, 37:1458–1465.

34. Stuart T, Butler A, Hoffman P, Hafemeister C, Papalexi E, Mauck WM, 3rd, Hao Y, Stoeckius M, Smibert P, Satija R: Comprehensive Integration of Single-Cell Data. Cell 2019, 177:1888–1902 e1821.

35. Romano S, Vinh NX, Bailey J, Verspoor K: Adjusting for chance clustering comparison measures. The Journal of Machine Learning Research 2016, 17:4635–4666.

36. Korsunsky I, Millard N, Fan J, Slowikowski K, Zhang F, Wei K, Baglaenko Y, Brenner M, Loh PR, Raychaudhuri S: Fast, sensitive and accurate integration of single-cell data with Harmony. Nat Methods 2019, 16:1289–1296.

37. Ma S, Zhang B, LaFave LM, Earl AS, Chiang Z, Hu Y, Ding J, Brack A, Kartha VK, Tay T, et al: Chromatin Potential Identified by Shared Single-Cell Profiling of RNA and Chromatin. Cell 2020, 183:1103–1116 e1120.

38. Liu J, Huang Y, Singh R, Vert JP, Noble WS: Jointly Embedding Multiple Single-Cell Omics Measurements. Algorithms Bioinform 2019, 143.

39. Welch JD, Kozareva V, Ferreira A, Vanderburg C, Martin C, Macosko EZ: Single-Cell Multiomic Integration Compares and Contrasts Features of Brain Cell Identity. Cell 2019, 177:1873–1887 e1817.

40. Hicks SC, Liu R, Ni Y, Purdom E, Risso D: mbkmeans: Fast clustering for single cell data using mini-batch k-means. PLoS Comput Biol 2021, 17:e1008625.

41. Shlyueva D, Stampfel G, Stark A: Transcriptional enhancers: from properties to genomewide predictions. Nat Rev Genet 2014, 15:272–286.

42. Pott S, Lieb JD: What are super-enhancers? Nat Genet 2015, 47:8–12.

43. Lieberman-Aiden E, van Berkum NL, Williams L, Imakaev M, Ragoczy T, Telling A, Amit I, Lajoie BR, Sabo PJ, Dorschner MO, et al: Comprehensive mapping of long-range interactions reveals folding principles of the human genome. Science 2009, 326:289–293.

44. Rao SS, Huntley MH, Durand NC, Stamenova EK, Bochkov ID, Robinson JT, Sanborn AL, Machol I, Omer AD, Lander ES, Aiden EL: A 3D map of the human genome at kilobase resolution reveals principles of chromatin looping. Cell 2014, 159:1665–1680.

45. Lock EF, Hoadley KA, Marron JS, Nobel AB: Joint and individual variation explained (JIVE) for integrated analysis of multiple data types. The annals of applied statistics 2013, 7:523.

46. Argelaguet R, Velten B, Arnol D, Dietrich S, Zenz T, Marioni JC, Buettner F, Huber W, Stegle O: Multi-Omics Factor Analysis-a framework for unsupervised integration of multiomics data sets. Mol Syst Biol 2018, 14:e8124.

47. Witten DM, Tibshirani R, Hastie T: A penalized matrix decomposition, with applications to sparse principal components and canonical correlation analysis. Biostatistics 2009, 10:515–534.

48. Shu H, Wang X, Zhu H: D-CCA: A decomposition-based canonical correlation analysis for high-dimensional datasets. Journal of the American Statistical Association 2020, 115:292–306.

