## Supplement for "Destin2: integrative and cross-modality analysis of single-cell chromatin accessibility data"

### *Supplementary Material*

**Supplementary Figure 1. UMAP visualization from unimodal analysis methods and Destin2's cross-modality analysis. Results are shown for the (A) PBMC, (B) adult mouse brain, (C) BMMC, and (D) human fetal tissue data.**

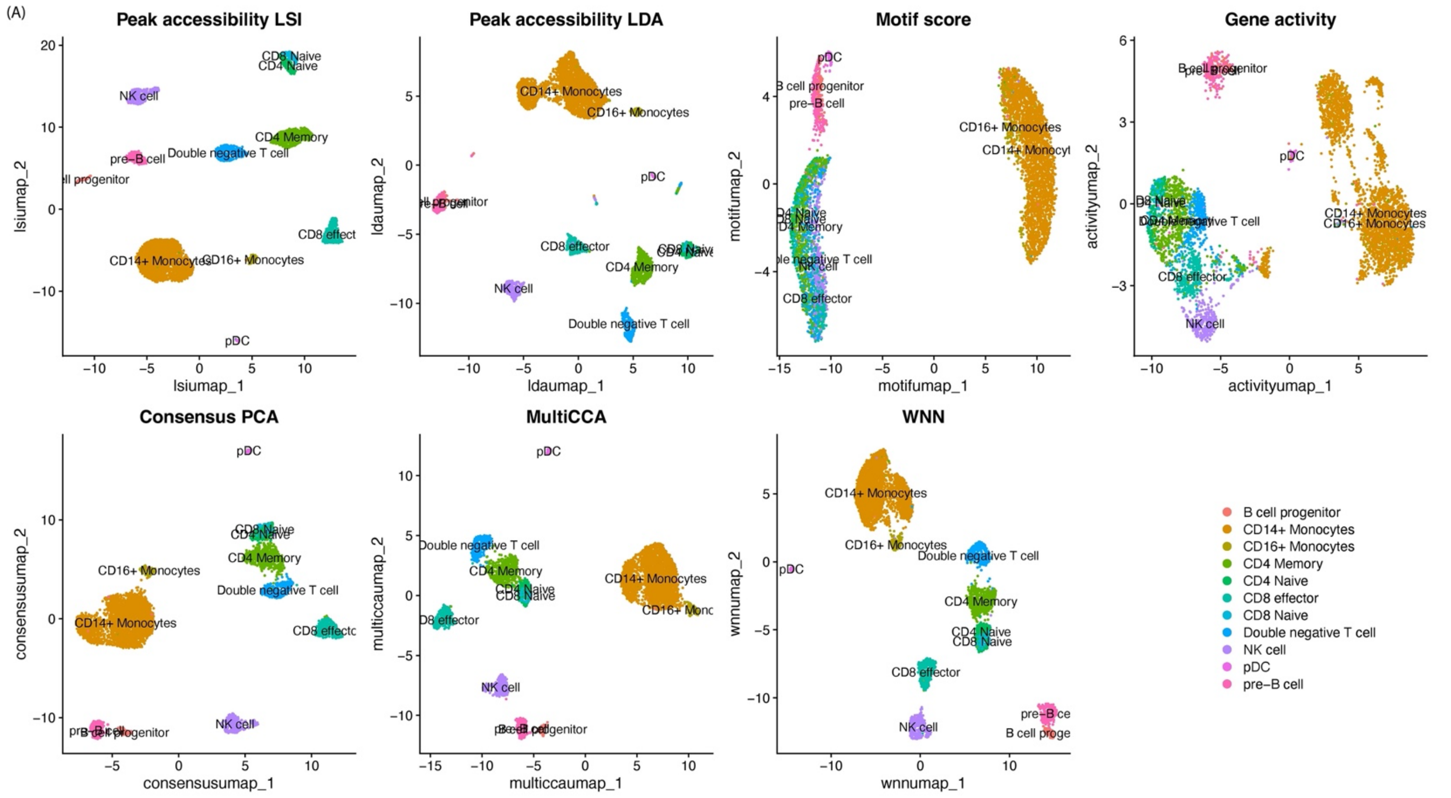

(B)

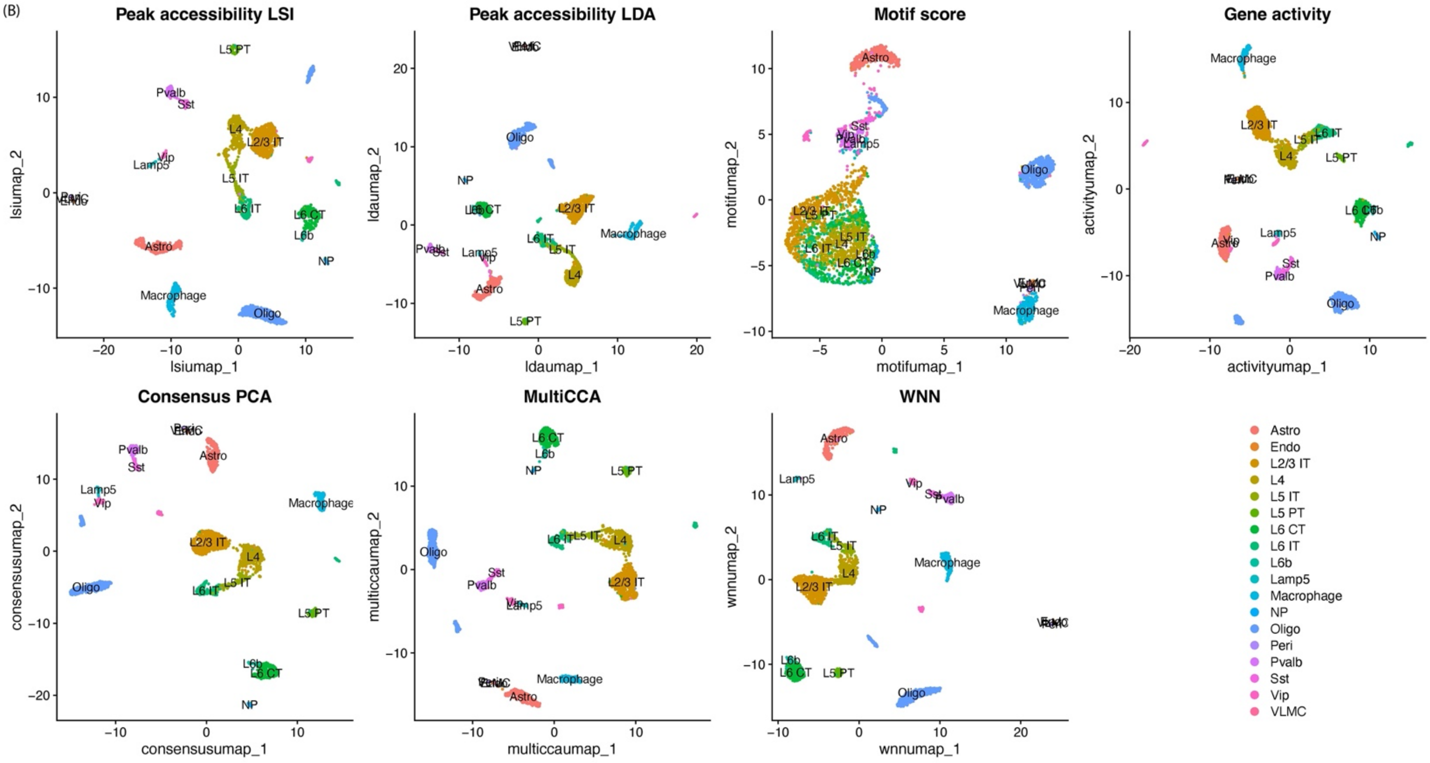
Peak accessibility LSI
lsiumap_1
lsiumap_2
Peak accessibility LDA
ldaumap_1
ldaumap_2
Motif score
motifumap_1
motifumap_2
Gene activity
activityumap_1
activityumap_2
Consensus PCA
consensusumap_1
consensusumap_2
MultiCCA
multiccaumap_1
multiccaumap_2
WNN
wnnumap_1
wnnumap_2
Astro
Endo
L2/3 IT
L4
L5 IT
L5 PT
L6 CT
L6 IT
L6b
Lamp5
Macrophage
NP
Oligo
Peri
Pvalb
Sst
Vip
VLMC

(C)

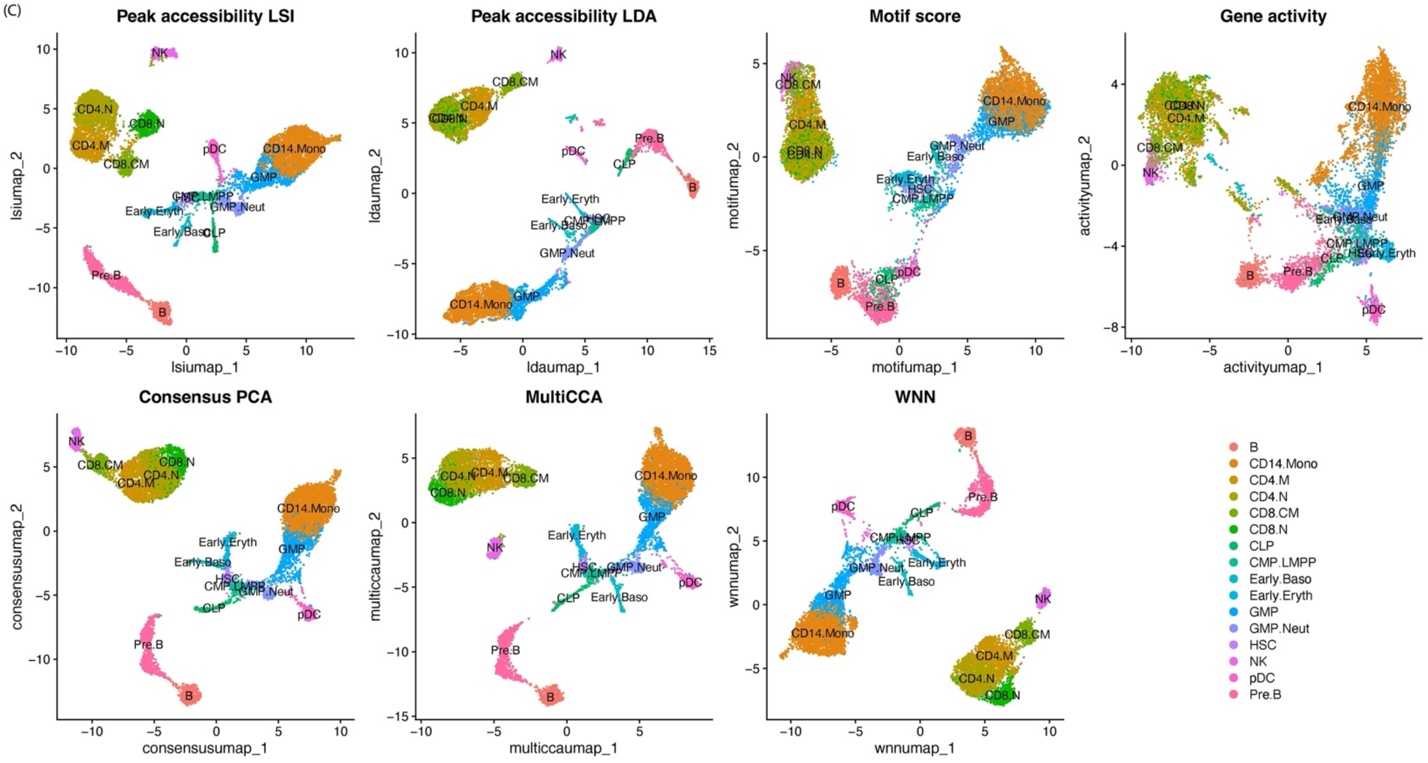
Peak accessibility LSI
lsiumap_1
lsiumap_2
Peak accessibility LDA
ldaumap_1
ldaumap_2
Motif score
motifumap_1
motifumap_2
Gene activity
activityumap_1
activityumap_2
Consensus PCA
consensusumap_1
consensusumap_2
MultiCCA
multiccaumap_1
multiccaumap_2
WNN
wnnumap_1
wnnumap_2
B
CD14.Mono
CD4.M
CD4.N
CD8.CM
CD8.N
CLP
CMP.LMPP
Early.Baso
Early.Eryth
GMP
GMP.Neut
HSC
NK
pDC
Pre.B

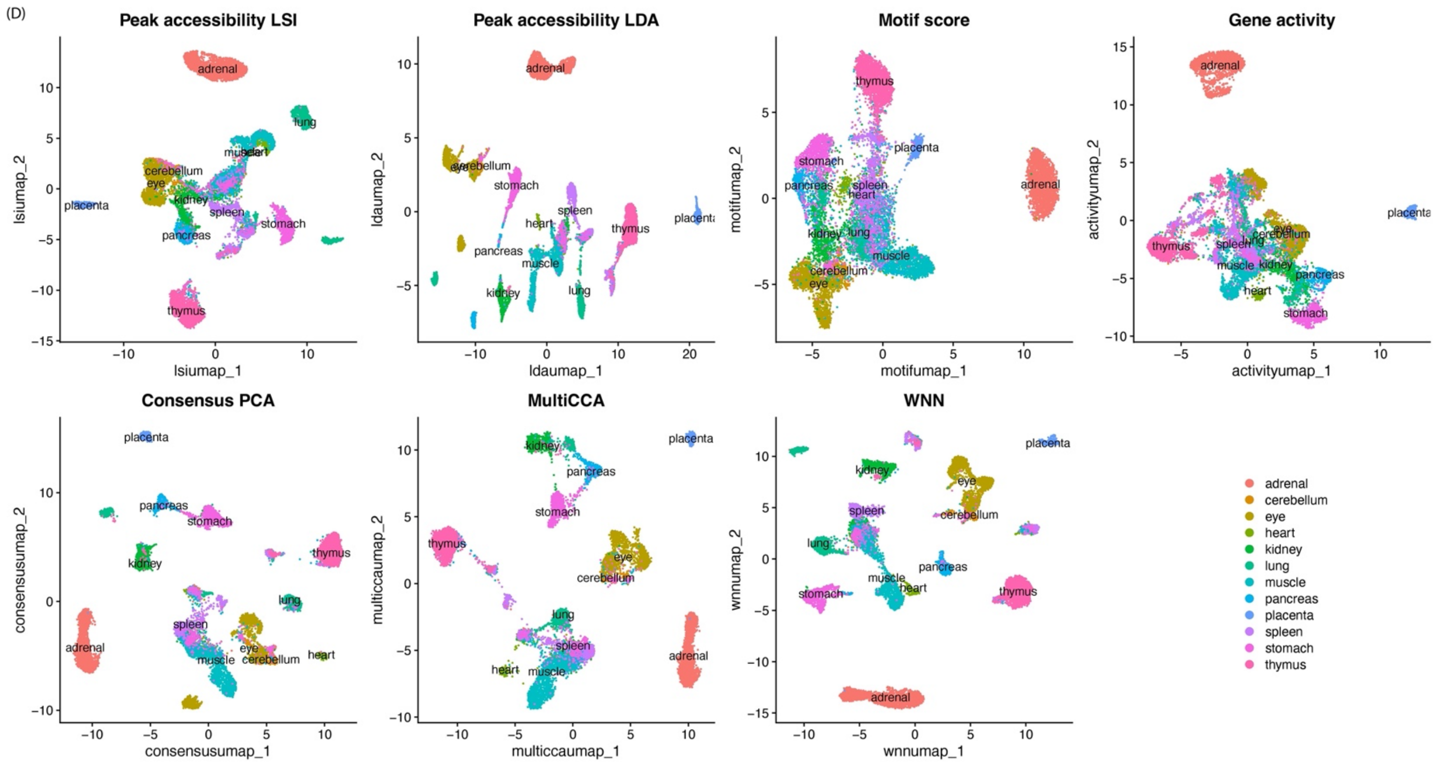


**Supplementary Figure 2.** Results with and without integrating the motif modality. Destin2 is robust to including a modality that does not contain as much information to separate the different cell types/states apart.

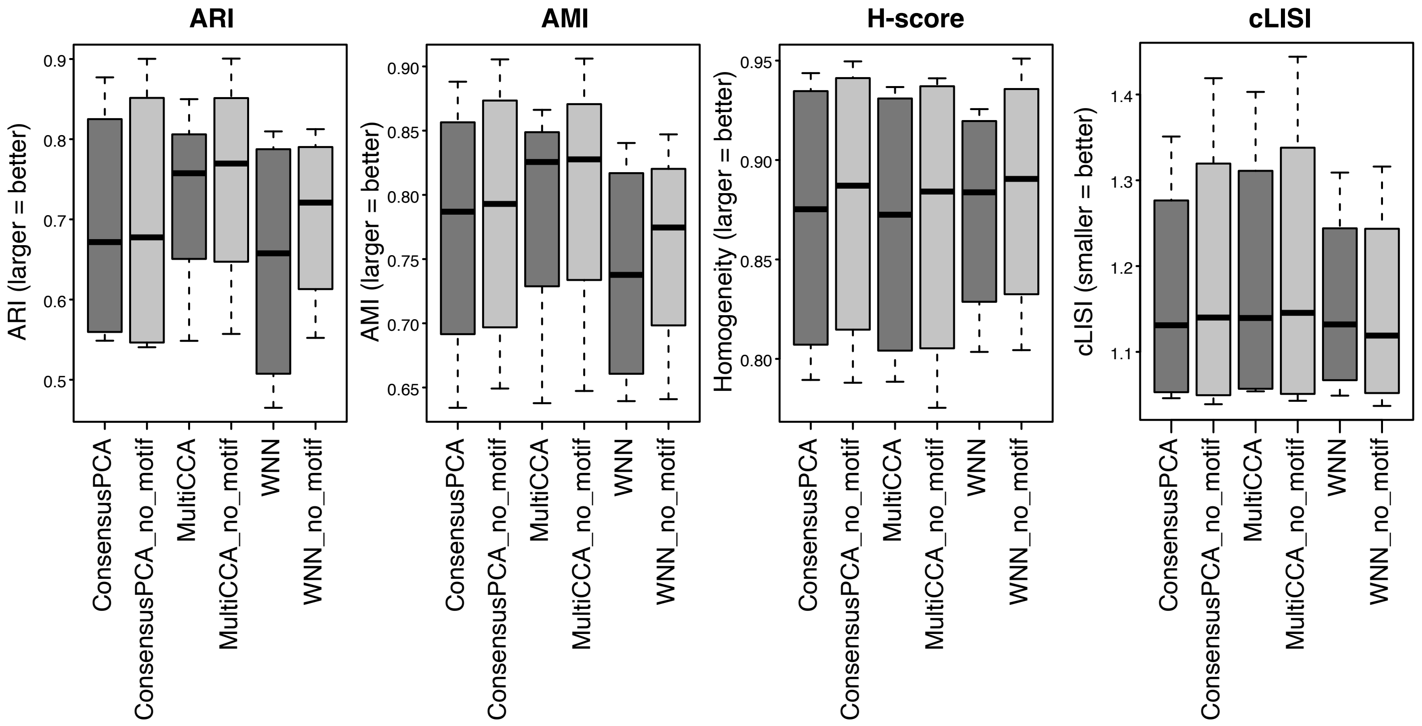

**Supplementary Figure 3.** Heatmap of confusion matrix. Cell types are transferred from single-cell RNA sequencing data; only cells with maximum prediction scores greater than 0.5 are kept in the analysis. Results are shown for the (A) PBMC, (B) adult mouse brain, (C) BMMC, and (D) human fetal tissue data.

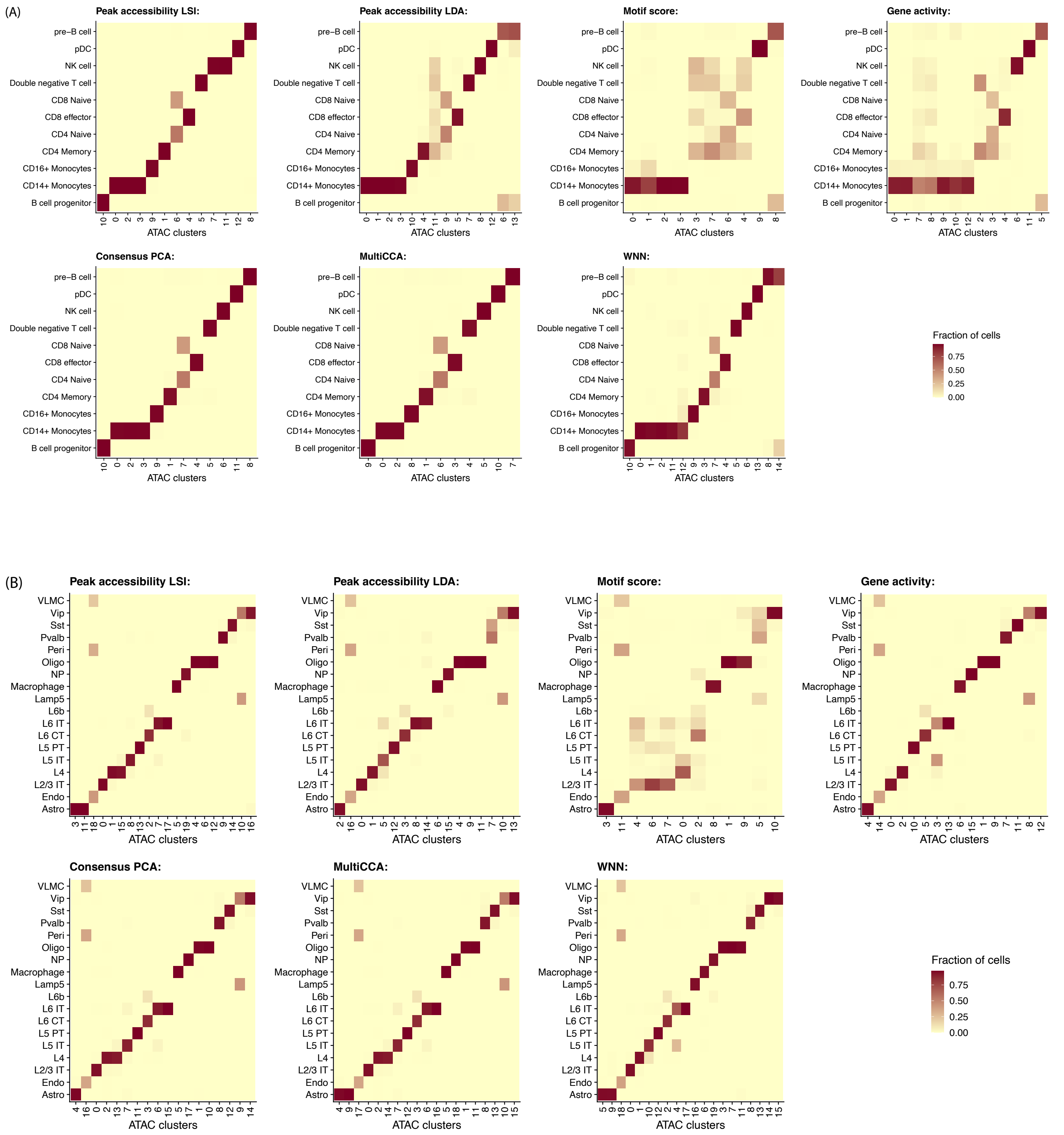

(C)

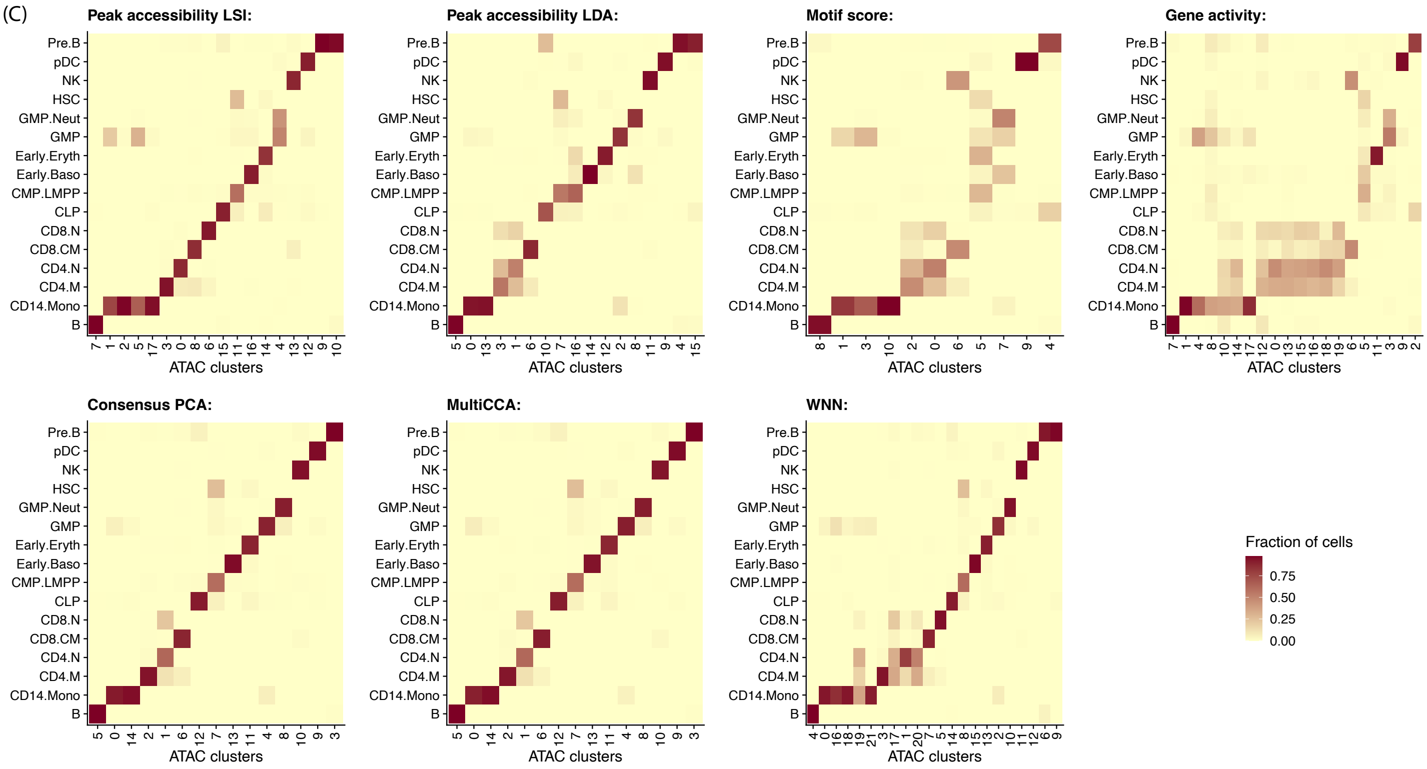
Peak accessibility LSI:
Peak accessibility LDA:
Motif score:
Gene activity:
Consensus PCA:
MultiCCA:
WNN:
Pre.B
pDC
NK
HSC
GMP.Neut
GMP
Early.Eryth
Early.Baso
CMP.LMPP
CLP
CD8.N
CD8.CM
CD4.N
CD4.M
CD14.Mono
B
ATAC clusters
Fraction of cells
0.75
0.50
0.25
0.00

(D)

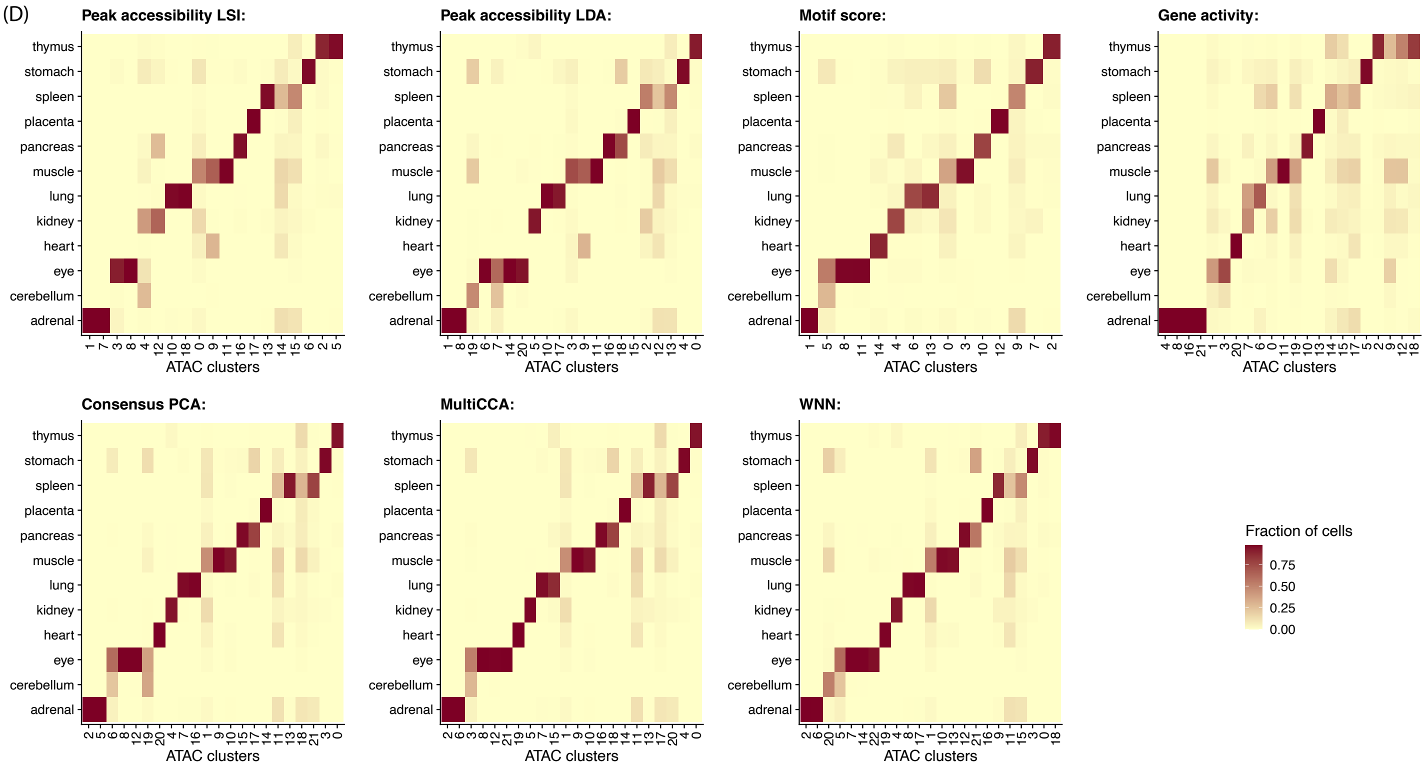
Peak accessibility LSI:
Peak accessibility LDA:
Motif score:
Gene activity:
Consensus PCA:
MultiCCA:
WNN:
thymus
stomach
spleen
placenta
pancreas
muscle
lung
kidney
heart
eye
cerebellum
adrenal
ATAC clusters
Fraction of cells
0.75
0.50
0.25
0.00

**Supplementary Figure 4.** Clustree output to determine the number of clusters. Results are shown for the (A) PBMC and (B) adult mouse brain data. The left and right panels show clustering trees with varying clustering resolutions and SC3 stability measures, respectively. New clusters form from existing clusters, and the overlap in cells between clusters at adjacent resolutions is computed and used to calculate the in-proportion for each edge. Unstable clusters result in cells switching between branches of the trees, with low in-proportion edges; one can thus infer which areas of the tree are likely to be the result of true clusters and which are caused by over-clustering.

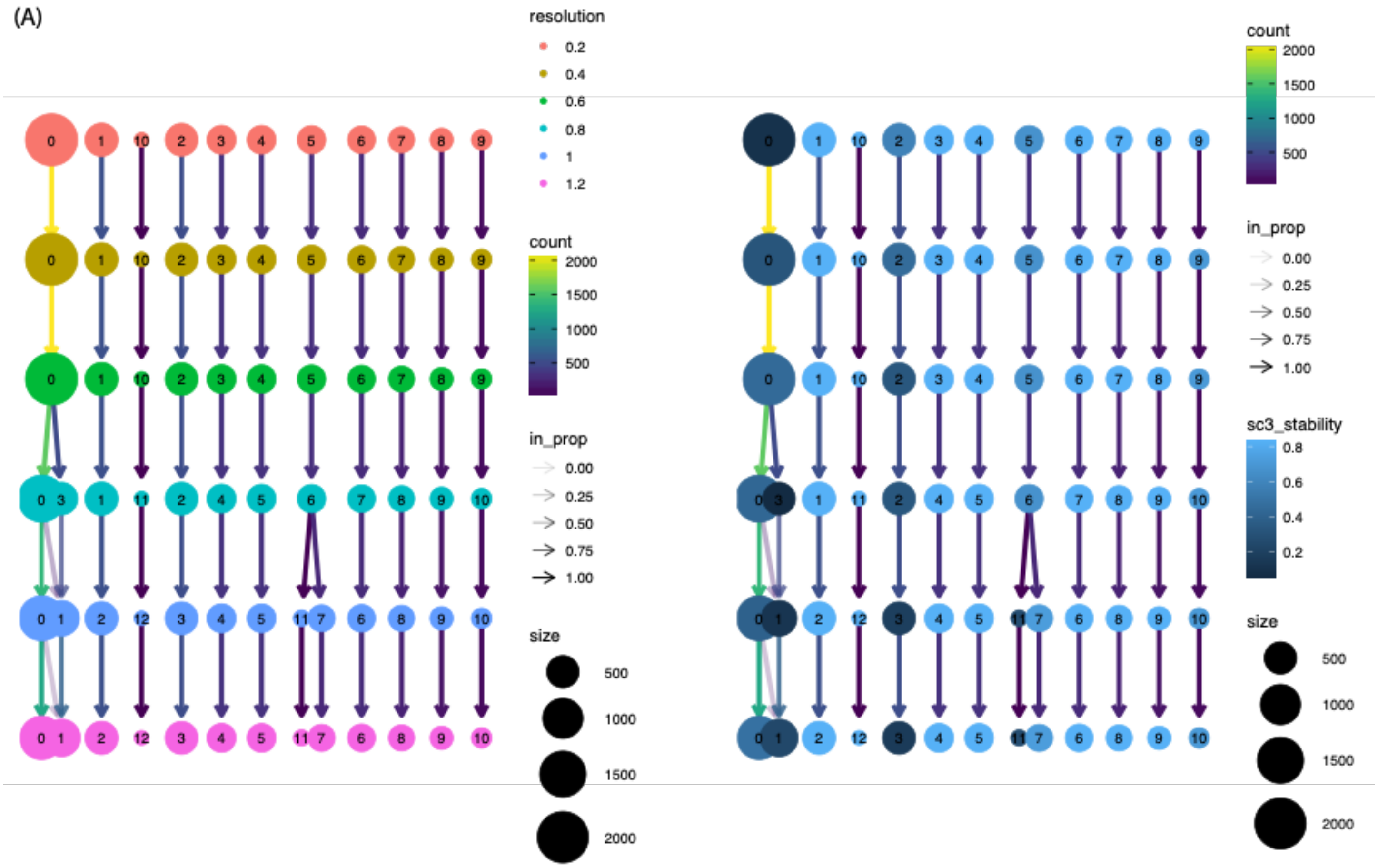

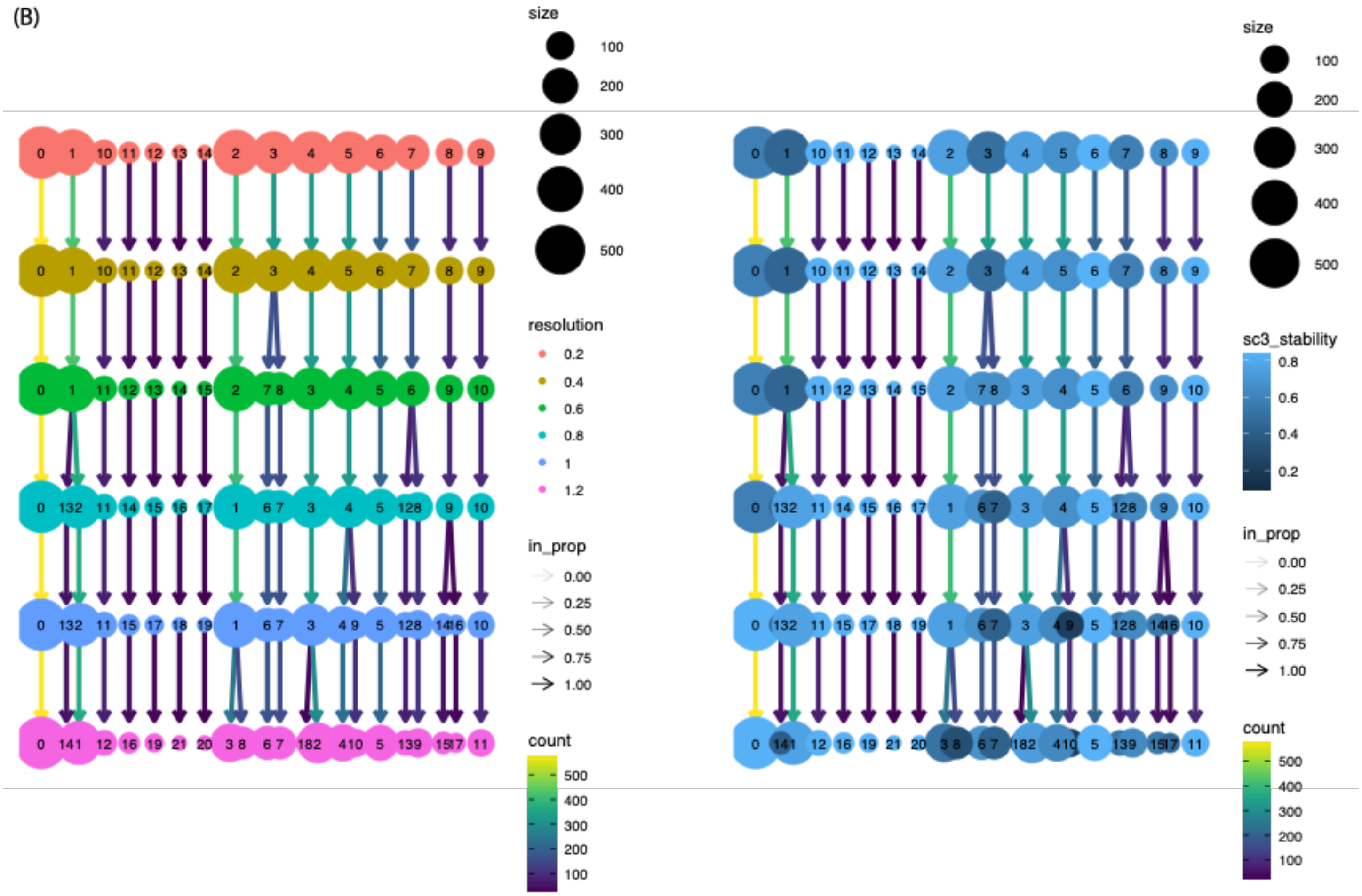


**Supplementary Figure 5.** Slingshot to reconstruct cell trajectory using Destin2's joint dimension reduction. Results are shown for the BMMC data of human hematopoietic differentiation.

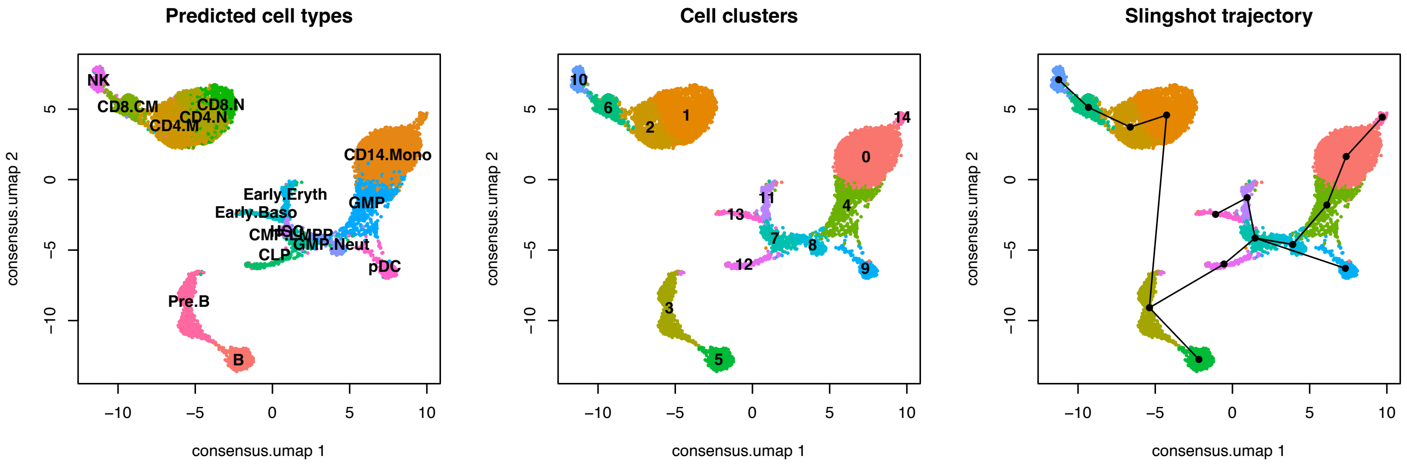

**Supplementary Table 1.** Data source and summary. Total number of cells and peaks, median number of ATAC reads per cell, median fraction of cells with detectable reads per peak, and number of annotated/transferred cell types are summarized post quality controls.

| Technology | Dataset | Source | # cells | # peaks | Median # reads per cell | Median frac cells per peak | # cell types / tissues |
|---|---|---|---|---|---|---|---|
| scATAC-seq | Human PBMC | 10x Genomics | 11017 | 80443 | 9333 | 0.041 | 16 |
| | Adult mouse cortex | 10x Genomics | 4972 | 87561 | 14600 | 0.023 | 11 |
| | Human BMMC | GSE139369 | 3196 | 154639 | 30121 | 0.044 | 18 |
| | Human fetal tissue | GSE149683 | 14275 | 1032191 | 3381 | 0.002 | 12 |
| scATAC+RNA multiomics | Human PBMC | 10x Genomics | 11331 | 108344 | 7306 | 0.021 | - |
| | Mouse embryonic brain | 10x Genomics | 4483 | 172150 | 8339 | 0.023 | - |
| | Mouse hair follicle | GSE140203 | 29308 | 343783 | 3364 | 0.005 | - |

**Supplementary Table 2.** Real data benchmarking results. Four different metrics – ARI, AMI, H-score, and cLISI – were used for performance assessment. Across the three scATAC-seq datasets and the four metrics, there is not a universally best method from the conventional unimodal analyses. On the other hand, Destin2's multimodal analyses achieve the best performance for ARI and AMI. For H-score and cLISI, since the LSI dimension reduction is used as weights in transferring the labels, it is not surprising that the peak LSI unimodal analysis achieves the best or near best performance. When all metrics are considered, Destin2 is the top rank method, as demonstrated in Figure 2.

| Data/Metric | | PBMC | | | | Mouse Brain | | | | BMMC | | | | Human fetal tissue | | | |
|---|---|---|---|---|---|---|---|---|---|---|---|---|---|---|---|---|---|
| | | ARI | AMI | H-score | cLISI | ARI | AMI | H-score | cLISI | ARI | AMI | H-score | cLISI | ARI | AMI | H-score | cLISI |
| Uni-modal | Peak (LSI) | 0.620 | 0.772 | 0.972 | 1.006 | 0.813 | 0.853 | 0.944 | 1.055 | 0.612 | 0.759 | 0.837 | 1.161 | 0.489 | 0.620 | 0.764 | 1.430 |
| | Peak (LDA) | 0.376 | 0.634 | 0.896 | 1.105 | 0.828 | 0.859 | 0.893 | 1.116 | 0.672 | 0.785 | 0.785 | 1.246 | 0.572 | 0.639 | 0.789 | 1.288 |
| | Motif | 0.282 | 0.467 | 0.589 | 1.645 | 0.559 | 0.663 | 0.663 | 1.709 | 0.469 | 0.633 | 0.633 | 1.629 | 0.588 | 0.648 | 0.696 | 1.729 |
| | Gene Activity | 0.473 | 0.570 | 0.684 | 1.325 | 0.860 | 0.891 | 0.891 | 1.091 | 0.502 | 0.622 | 0.622 | 1.830 | 0.358 | 0.478 | 0.591 | 1.691 |
| Multi-modal | CPCA | 0.570 | 0.749 | 0.944 | 1.046 | 0.877 | 0.888 | 0.926 | 1.060 | 0.773 | 0.825 | 0.825 | 1.202 | 0.549 | 0.634 | 0.789 | 1.351 |
| | MultiCCA | 0.753 | 0.831 | 0.937 | 1.054 | 0.850 | 0.866 | 0.925 | 1.060 | 0.762 | 0.820 | 0.820 | 1.219 | 0.549 | 0.638 | 0.789 | 1.403 |
| | WNN | 0.465 | 0.682 | 0.926 | 1.049 | 0.810 | 0.840 | 0.914 | 1.085 | 0.765 | 0.793 | 0.854 | 1.179 | 0.550 | 0.639 | 0.804 | 1.309 |
